# Pangenome reference assemblies reveal the variation and recent activity of human LINE-1 retrotransposons

**DOI:** 10.64898/2026.05.14.725010

**Authors:** Lei Yang, Sara Nematbakhsh, Amanda Norseen, Richard N. McLaughlin

## Abstract

LINE-1 retrotransposons are the only autonomous mobile elements still active in human genomes and remain a potent source of mutation, genome remodeling, and disease risk. However, young, full-length, potentially active copies (the elements most likely to shape present-day genomes) have been largely inaccessible to population-scale analysis because they are long, repetitive, and poorly resolved by short-read sequencing. Here, we use 47 phased long-read assemblies from the Human Pangenome Reference Consortium, representing 94 haplotypes, to build an allele-resolved view of recent human LINE-1 evolution. We identify 13,617 LINE-1 alleles with intact ORF1 and ORF2 across 683 unique insertion sites, revealing that every genome carries a distinct repertoire of potentially active source elements. These intact LINE-1 profiles recapitulate broad human population structure while exposing a large, rare, and population-enriched reservoir of mobile-element diversity missed by single-reference approaches. We also resolve a structurally variable chromosome 11 LINE-1 array, demonstrating that local duplication and rearrangement can amplify LINE-1 sequence independently of canonical retrotransposition. By comparing full-length LINE-1 sequences, we define activity signatures that separate ancient remnants from recently expanding lineages and uncover young LINE-1 groups whose activity is not fully explained by canonical subfamily labels. Sequence-network analyses further reveal a dynamic history of lineage turnover, in which successful source elements rise, seed new insertions, and are replaced by descendants marked by specific nucleotide changes. Together, these data transform human LINE-1s from a repetitive background into a resolved evolutionary system, linking insertion polymorphism, coding potential, population history, and recent retrotransposon adaptation. Our findings establish the human pangenome as a framework for discovering active source elements and for testing how mobile DNA continues to shape genome evolution, host defense, and disease risk.

## Introduction

Transposable elements have shaped human genome evolution through repeated cycles of proliferation, host restriction, and molecular decay [1,2]. Although most transposable element copies in the human genome are no longer mobile, their sequences continue to influence genome architecture, gene regulation, chromatin state, and structural variation [3,4]. Among extant human transposable elements, LINE-1 elements are unique because they remain the only autonomous retrotransposons with measurable activity [5,5]. Full-length LINE-1s encode the proteins required for their own mobilization and can also mobilize non-autonomous elements, including Alu and SVA, as well as processed host transcripts [6–8]. As a result, LINE-1 activity remains a continuing source of human genetic variation [9,10].

The mutagenic potential of LINE-1 retrotransposition has selected for multiple layers of host control. DNA methylation, chromatin-based repression, RNA surveillance, and post-transcriptional restriction factors all act to limit LINE-1 expression or mobilization [11]. These defenses are expected to impose evolutionary pressure on active LINE-1 lineages, while successful LINE-1 variants may in turn acquire sequence changes that alter expression, protein function, RNA processing, or evasion of host restriction [11,12]. Human LINE-1 evolution therefore reflects both the population history of insertions and an ongoing molecular conflict between retrotransposon activity and host defense.

LINE-1 insertions can have large functional consequences despite being relatively rare as new germline events. De novo mobile element insertions occur in roughly one of every 20-40 births across mobile element classes, and LINE-1-mediated insertions have been linked to inherited disease, cancer, and somatic mosaicism [4,10,13,14]. However, most prior population-scale studies of human LINE-1 variation have focused on insertion presence or absence [15–18]. The sequence-level diversity of full-length LINE-1 alleles has remained more difficult to resolve, even though these allelic differences are likely central to retrotransposition potential, source-element activity, and interactions with host restriction factors [19–21].

This gap has been largely technical: LINE-1 elements are approximately 6 kb long and occur in hundreds of thousands of copies across the genome, with thousands of full-length copies embedded in repetitive sequence [1,22]. Short-read sequencing often cannot span an entire element together with enough unique flanking sequence to place the insertion confidently or reconstruct allelic sequence differences. Consequently, reference-based and short-read approaches have likely underestimated both the diversity of full-length LINE-1 insertions and the number of potentially active source elements segregating in human populations [16–18].

Long-read, haplotype-resolved genome assemblies now make it possible to study LINE-1 variation at a more appropriate scale. The Human Pangenome Reference Consortium provides diverse phased assemblies that capture both reference and non-reference insertions, including alleles that are rare, population-enriched, or absent from any single reference genome [23]. This framework enables LINE-1 insertions to be compared across haplotypes, assigned to shared insertion sites, and analyzed not only for presence or absence but also for intactness and full-length sequence variation [21,24].

Resolving this recent phase of LINE-1 evolution is especially relevant because the finest evolutionary signals are transient. Shortly after insertion, related LINE-1 copies retain the sequence similarity, intact ORFs, flanking transduction signatures, and population-frequency patterns that connect mothers to daughters [25–27]. Over longer timescales, mutation obscures these relationships, leaving older LINE-1 families informative for broad subfamily history but less able to reveal the fine-scale dynamics of source turnover [12,28]. Young, full-length LINE-1s in haplotype-resolved pangenomes therefore provide an unusually detailed record of retrotransposon evolution in action.

Here, we use phase-1 HPRC assemblies to characterize human LINE-1 diversity, recent activity, and sequence evolution across 47 individuals and 94 haplotypes [23]. We identify 13,617 LINE-1 alleles with intact ORF1 and ORF2 across 683 unique insertion sites, revealing that each individual carries a distinct repertoire of intact LINE-1 alleles. We then use allele-resolved presence, absence, and intactness calls to describe how these elements are distributed across human populations and to estimate the broader landscape of intact LINE-1 diversity beyond the sampled cohort.

Finally, we use full-length LINE-1 sequence variation to infer recent activity and evolutionary turnover. Neighbor-distance profiles identify groups of LINE-1s with signatures of recent *in vivo* expansion, while sequence-network analyses reveal transitions among young LINE-1 lineages and nominate nucleotide changes associated with recent diversification. Together, these analyses show how pangenome assemblies can connect LINE-1 burden, population distribution, recent source activity, and adaptive sequence change in a single allele-resolved framework.

## Results

### HPRC assemblies enable an allele-resolved catalog of full-length and intact LINE-1s

We analyzed phase-1 Human Pangenome Reference Consortium assemblies comprising 47 individuals and 94 phased haplotypes. Because these assemblies are long-read and haplotype-resolved, they present a resource that is uniquely poised to study LINE-1 variation. Specifically, with average sequencing read lengths longer than the 6,000 bp LINE-1 repeat unit, the full internal sequence of individual elements can be recovered along with their flanking sequence, assigned to homologous insertion sites, and compared across haplotypes. This pangenome-scale approach extends beyond deep, single-reference genomes and more-shallow population scale views of human genome structure and repeat annotation, including the T2T-CHM13/hs1 reference, the first HPRC assemblies, and related resources [23,24,29,30].

We identified full-length LINE-1 sequences in each haplotype assembly using RepeatMasker [31] annotations and the HapLongLINEr pipeline (https://github.com/leiyangly/HapLongLINEr), which checks each element for intact ORF1 and ORF2 (see methods), and grouped alleles from different haplotypes into homologous/shared insertion sites by lifting flanking sequence to hs1. Each haplotype assembly contained thousands of full-length LINE-1s, providing a genome-by-genome inventory of LINE-1 content. At the pangenome/population level, we resolved 2,275 recent full-length LINE-1 insertion sites, including 683 sites where at least one haplotype carried an intact allele. Within the population of all 94 haplotypes, these intact-LINE-1 containing sites accounted for 13,617 intact LINE-1 alleles.

At the haplotype level, HPRC assemblies carried 9,251 ± 320 full-length LINE-1s on average, with 145 ± 9 intact LINE-1s per haplotype (Figure 1A, B). The largest systematic difference in LINE-1 abundance among haplotypes reflected sex-chromosome content: X-bearing haplotypes carried more full-length LINE-1s than Y-bearing haplotypes on average and also carried a modestly higher number of intact LINE-1s. At the diploid level, individuals carried 18,502 ± 400 full-length LINE-1s and 290 ± 10 intact LINE-1s on average (Figure 1C, D). XX genomes carried more full-length and intact LINE-1s than XY genomes, consistent with the larger repeat content of chromosome X relative to chromosome Y. These diploid intact counts are consistent with the approximately 300 intact LINE-1s calculated from combining the CHM1 and CHM13 haploid references, supporting the consistency of the HPRC call set with prior long-read, allele-resolved LINE-1 analyses [21].

**Figure 1.**
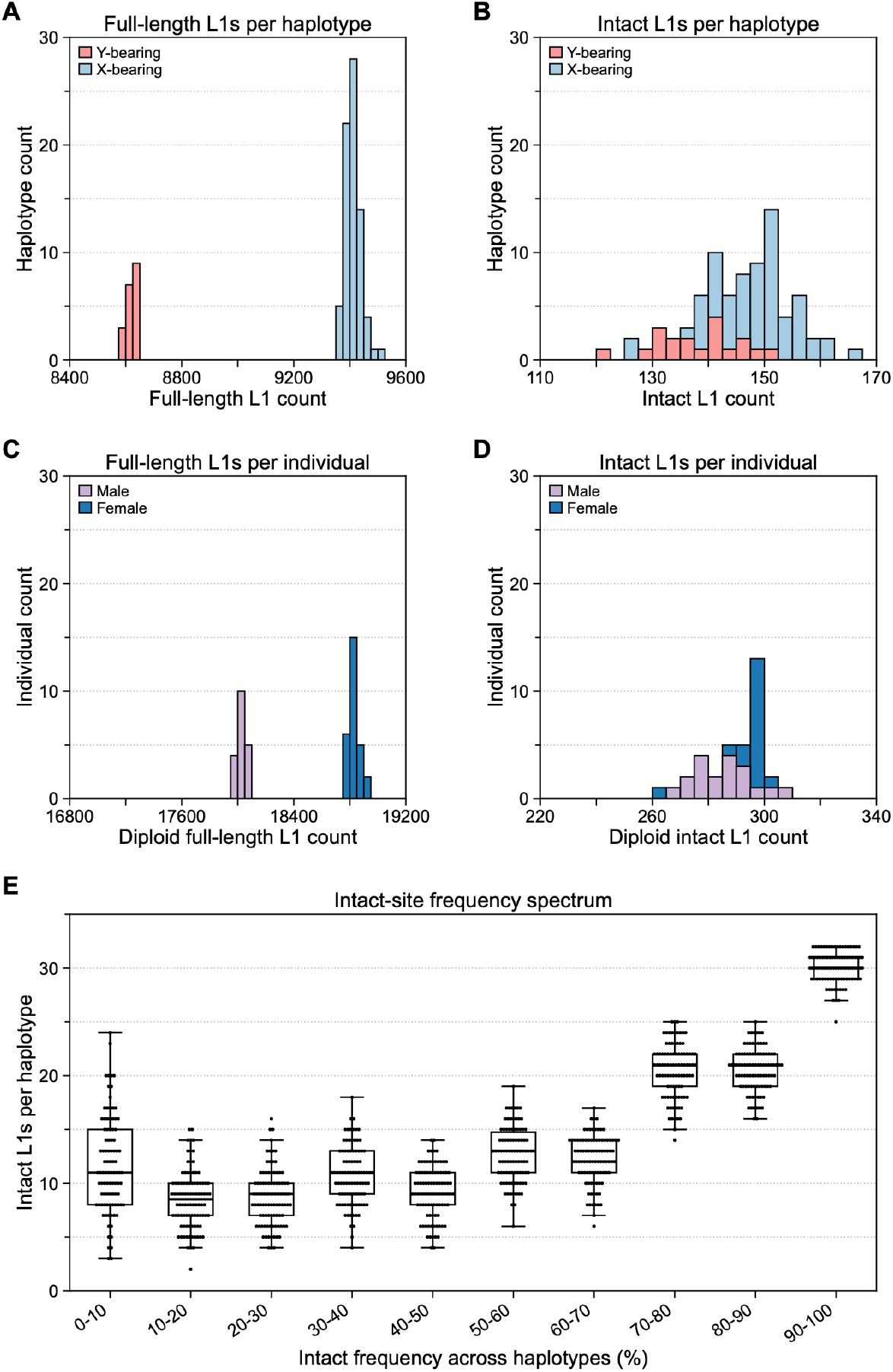
Full-length and intact LINE-1 burden across HPRC haplotypes and individuals. (A) Distribution of full-length LINE-1 counts per haplotype across 94 phased HPRC haplotypes. X-bearing haplotypes are shown in light blue and Y-bearing haplotypes in pink. (B) Distribution of intact LINE-1 counts per haplotype across the same haplotypes. (C) Distribution of diploid full-length LINE-1 counts per individual across 47 HPRC individuals. Female genomes are shown in blue and male genomes in lavender. (D) Distribution of diploid intact LINE-1 counts per individual. (E) Distribution of intact LINE-1 loci across the intact-frequency bins of all haplotypes. For each intact LINE-1 insertion site, cohort-wide intact frequency was calculated as the fraction of eligible HPRC haplotypes carrying an intact allele at that site, using chromosome-aware denominators: 94 haplotypes for autosomal loci, 75 X-bearing haplotypes for chromosome X loci, and 19 Y-bearing haplotypes for chromosome Y loci. Sites were assigned to 10 frequency bins from 0-10% through 90-100%. Each bin contains one point per haplotype (n = 94), representing the number of intact LINE-1 loci in that haplotype whose cohort-wide intact frequency falls within the indicated frequency interval. Boxplots summarize the distribution across haplotypes; overlaid points show individual haplotypes.

As previously described [21], phasing and grouping syntenic alleles allowed us to classify all insertion sites in each haplotype into one of three allele states: (1) absent, (2) present but interrupted by SNPs or indels that disrupt ORF1 and/or ORF2, or (3) intact, retaining both ORF1 and ORF2 coding potential. In diploid genomes, this resolves loci that are intact on both haplotypes from loci where an intact allele is paired with an interrupted allele or with absence of the insertion on the homologous haplotype. This tri-state framework captures information lost in presence/absence catalogs from earlier population-scale MEI surveys [15,17,18]. This classification allowed us to simultaneously analyze LINE-1 variation as population variation, molecular-state variation, and potential source-element diversity.

### Individual haplotypes contain both common and low-frequency intact LINE-1 alleles

We next asked how much of each haplotype’s intact LINE-1 repertoire is shared broadly across the HPRC cohort and how much lies in the low-frequency tail of segregating variation. Figure 1E summarizes how intact LINE-1 alleles are distributed across cohort frequency bins within each haplotype. Each haplotype contains a shared high-frequency core of intact loci, with median counts of 21 intact LINE-1s in the 70–80% bin, 21 in the 80–90% bin, and 30 in the 90–100% bin. However, lower-frequency bins remain consistently populated, with median counts ranging from 8.5 to 13 intact loci per bin across the 0–10% through 60–70% ranges. Thus, human haplotypes are not simply a collection of fixed or near-fixed intact LINE-1s. Instead, each haplotype contains a high-frequency core of intact alleles shared by many individuals and a broad tail of low- and intermediate-frequency intact alleles. The high-frequency core may represent broadly shared source-element candidates that contribute to baseline human LINE-1 activity, whereas the lower-frequency tail is enriched for alleles that are more recent, more population-structured, or less completely sampled. These data reinforce that frequency alone cannot distinguish age from activity.

### LINE-1 allele-state profiles recapitulate human population structure

To visualize how intact LINE-1 alleles are distributed across people, we encoded each intact insertion site in each haplotype as absent, present but interrupted, or intact, and plotted these states across the HPRC cohort (Figure 2). The autosomal panel includes 648 loci across all 94 haplotypes, while the sex-chromosome panel displays 34 X-linked loci and one Y-linked locus separately across 75 X-bearing and 19 Y-bearing haplotypes. The 34 X-linked loci and single Y-linked locus did not show a detectable population-associated distribution pattern (Figure 2B). The autosomal ordering recovers a strong population axis where most African haplotypes form a broad block, whereas admixed American, East Asian, European, and South Asian haplotypes have an expected intermediate structure, consistent with prior mobile-element and genome-wide variant analyses showing that insertion and SNP profiles track human ancestry [17,32].

**Figure 2.**
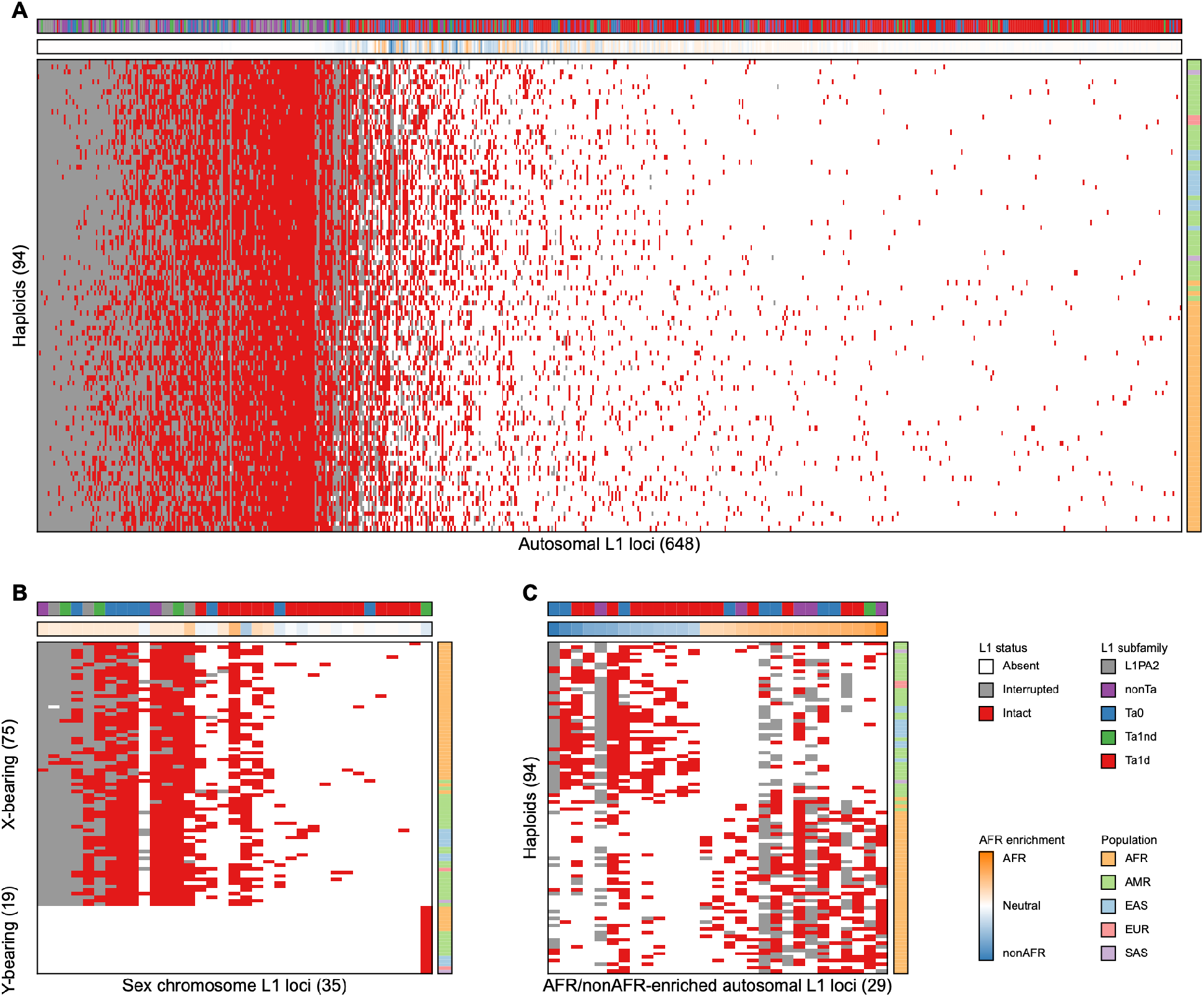
Population distribution of intact and interrupted LINE-1 alleles across HPRC haplotypes. (A) Autosomal LINE-1 insertion sites carrying an intact allele in at least one haplotype, shown across 94 phased HPRC haplotypes. Cells indicate absence, present but interrupted, or intact LINE-1 status; top annotation bars show AFR enrichment and LINE-1 subfamily, and the side bar shows 1000 Genomes super-population assignment. (B) Sex-chromosome LINE-1 sites shown with the same status encoding, with X-bearing and Y-bearing haplotypes separated on the y-axis. (C) Autosomal LINE-1 sites significantly enriched between AFR and non-AFR haplotypes. Enrichment is shown as the difference in presence frequency between AFR and non-AFR haplotypes, and significance was defined after multiple-testing correction. Together, these panels show that LINE-1 presence and intactness recapitulate broad population structure while retaining a substantial rare and population-enriched tail.

LINE-1 patterns within admixed American haplotypes are consistent with known ancestry structure. For example, Peru-associated haplotypes trend toward the East Asian side of the non-African axis, reflecting the deep relationship between Native American and East Asian lineages [33,34]. Colombian haplotypes from Medellín fall closer to Europeans, matching prior analyses showing predominantly European ancestry in that population, with lower Native American and African contributions on average [35,36].

### LINE-1 presence and intactness show patterns of population enrichment

Within the autosomal intact set, 196 LINE-1 insertion sites were observed only in African haplotypes and 82 were observed only in non-African haplotypes, but population specificity was more broadly apparent as enrichment rather than strictly private insertions. We analyzed the relative abundance of autosomal insertions between African and non-African haplotypes and identified 29 loci with significant AFR/non-AFR enrichment at FDR 5%, including 16 AFR-enriched and 13 non-AFR-enriched LINE-1 insertions (Figure 2C). This pattern is consistent with prior MEI catalogs in which most insertions are shared across populations but differ substantially in frequency, especially among rare and non-reference LINE-1s [17,18,37].

Because the HPRC panel is enriched for African and admixed American genomes but has limited European and South Asian representation, these contrasts are best interpreted as broad AFR/non-AFR signals rather than a complete fine-scale map of continental specificity of LINE-1s. Nevertheless, the enrichment analysis provides a quantitative summary of the population structure visible in Figure 2A and shows that intact LINE-1 insertion sites are present at different allele frequencies across human populations.

### Intactness profiles reflect the temporal trajectory of LINE-1 insertion and decay

The tri-state intactness view of the HPRC haplotypes also reinforces the basic temporal view of LINE-1 evolution. LINE-1s retrotranspose to create newly inserted LINE-1s. These singleton LINE-1s can be either intact or (due to the mutagenic- and 5’ truncating-nature of LINE-1 reverse transcriptase) present but interrupted. Over time, some intact singleton LINE-1s increase in frequency, while many accumulate internal mutations, including SNPs, indels, and rearrangements that convert them into present-but-interrupted alleles, a pattern consistent with prior analyses of LINE-1 insertion structure and retrotransposition intermediates [38,39].

Consistent with this model, 155 autosomal intact loci are present in all 94 haplotypes, but only two are intact in all haplotypes. This suggests the importance (especially in high frequency LINE-1s) of differentiating how often an insertion is present and how often it remains intact among carriers. Some LINE-1 insertions are present in nearly all haplotypes but retain intact ORF1 and ORF2 in only a subset, indicating that the insertion itself is old while intact coding potential has eroded probabilistically on some haplotypic backgrounds. In contrast, LINE-1s that are high frequency and frequently intact define a shared core of preserved source-element candidates that have been inherited and preserved from ancestral haplotypes.

It follows that high-frequency LINE-1s include both older insertion sites undergoing molecular decay and broadly shared intact alleles that have retained coding potential. This contrast suggests a temporal model in which LINE-1 insertions enter populations as intact alleles, then follow different trajectories: loss, persistence as rare intact variation, expansion to common intact alleles, or decay into common interrupted alleles. In addition, each genome contains a set of broadly shared intact LINE-1s that may have already been present in the last common ancestor of the sampled haplotypes and have retained ORF1 and ORF2 intactness across many generations. Thus, the intact LINE-1 repertoire of any individual reflects both recent, low-frequency variation and an older shared core of preserved source-element candidates. This detailed population distribution can be leveraged into an evolutionary signal – insertion presence reflects ancestry and drift, intactness represents putative retrotransposition potential, and the combination of the two reveals how LINE-1 alleles move from rare intact copies toward older interrupted forms.

### Tandem duplicated LINE-1s at chromosome 11 reveal local structural complexity in LINE-1 evolution

The analyses above treat LINE-1s as retrotransposition-defined, homoplasy-free insertion sites that can be grouped across haplotypes by shared flanking sequence. However, our pipeline identified an unusually large number of LINE-1 annotations mapping to a single locus on chromosome 11 (hs1, ch11:94,140,954-94,254,695). Closer inspection showed that this signal did not reflect many independent retrotransposition events at the same site, but instead a complex LINE-1-rich region on chromosome 11 that contains 16 back-to-back young, L1HS LINE-1 copies in the hs1 reference. In hg38, this region contains no resolved L1HS sequences, likely due to a break in the assembly.

The chromosome 11 locus is bounded by conserved unique flanking sequence in addition to older LINE-1 sequence on either side of the variable region. These conserved flanks allowed the locus to be located and aligned across haplotypes and distinguished the internal hypervariable region from the surrounding genomic context. Similar to recent studies of structurally complex repetitive regions [40], this approach allowed us to examine both the organization of the repeat array and the sequence relationships among individual LINE1 copies across diverse human haplotypes. Within the variable interval, LINE-1-derived units are arranged as a tandem array rather than as isolated insertions. Thus, this locus likely represents the local expansion and contraction of a LINE-1-containing array, rather than the canonical presence or absence of a single retrotransposed element.

We next examined the composition of the array across HPRC haplotypes. Upon further examination of this region in the HPRC assemblies, we find a highly variable local structure in which the number of LINE-1 units ranged from 0 to 15 across haplotypes (Figure 3). Although the LINE-1 units in this array are not intact by the ORF1/ORF2 definition used above, many are longer than 5,000 bp and retain a 5′UTR and intact ORF1p-coding region. Individual haplotypes differed not only in total LINE-1 copy number, but also in the order and identity of LINE-1 units within the locus. Sequence comparison of the array units revealed substantial heterogeneity, indicating that copies within the locus are not interchangeable repeats of a single identical element. Instead, the repeat units fall into multiple sequence groups. Despite this variation, two core repeat units seem to account for much of the structural diversity at the locus, suggesting that recurrent duplication, deletion, or rearrangement of these units generated many of the observed haplotypic configurations.

**Figure 3.**
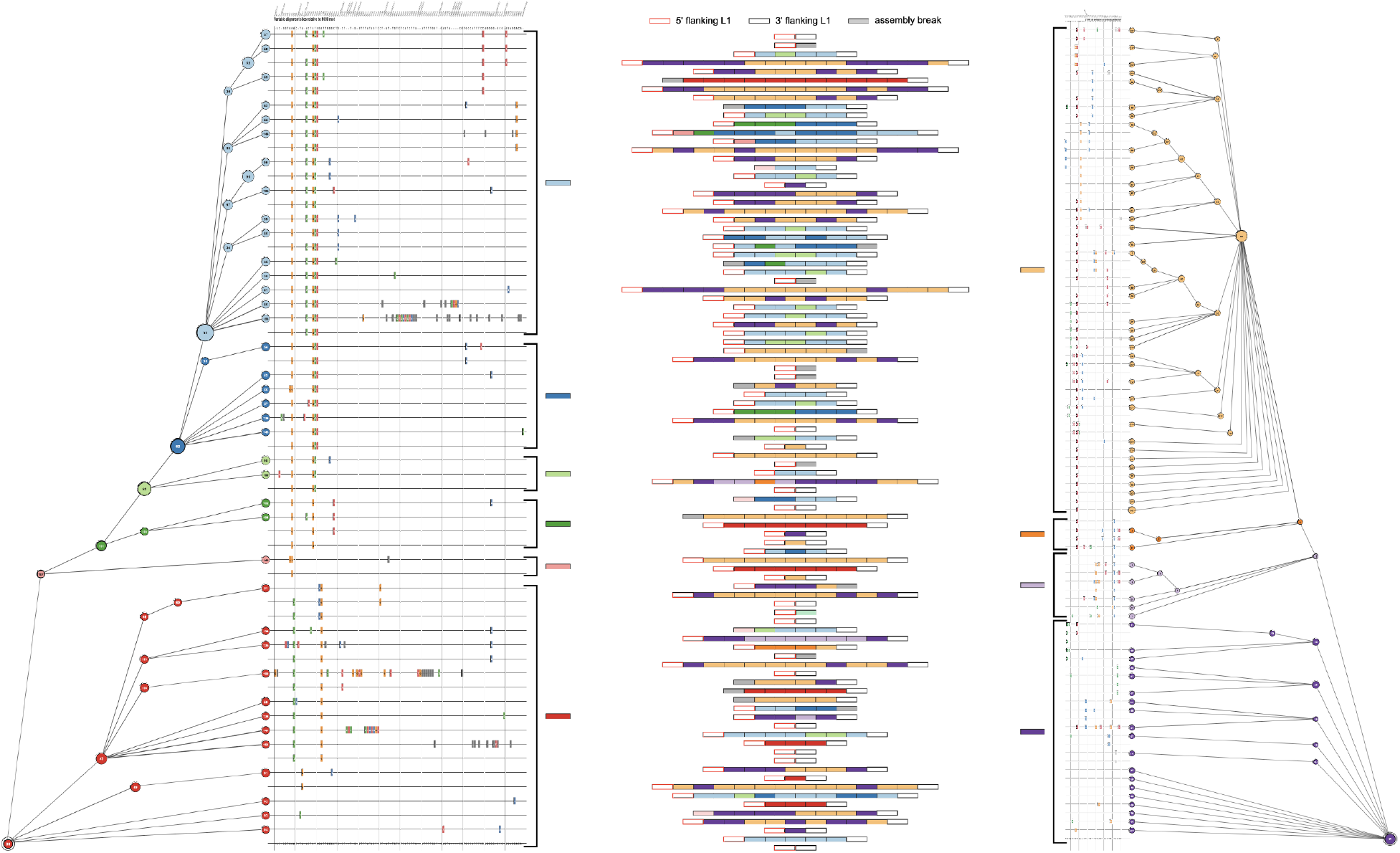
Structure of a polymorphic chr11 tandemly-duplicated LINE-1 array across HPRC haplotypes. Left, right: A minimum-spanning network and alignment of variable positions from full-length L1HS copies (>5600 bp) from the chr11 LINE-1 array. Each numbered node represents a unique L1HS sequence; identical L1HS copies from different haplotypes were collapsed into a single node, and node size is proportional to the number of copies assigned to that variant. The network topology showed two clusters, which we split. N090 and N072 were used as the root/reference sequence for the two clusters and adjacent alignment matrix. Rows in the alignment correspond directly to numbered nodes in the network, and columns show variable nucleotide positions relative to the root after removing singleton sites. Colored cells indicate bases that differ from root. Node colors indicate major backbone lineages (N031, N052, N095, N101, N107, N090, N106, N003, N011, and N072), with descendant nodes assigned the same color as their corresponding backbone node. Middle: Haplotype-array heatmap showing the structure and sequence relationships of chr11 LINE-1 copies across HPRC haplotypes. Each row represents one haplotype, and columns represent LINE-1 copies ordered by genomic position within the chr11 array. Tile colors match the node/lineage colors used in the networks, linking each array copy to its corresponding sequence variant or lineage. The 5’ flanking LINE-1 (L1PA2) and 3’ flanking LINE-1 (L1P1) copies are shown as empty red or black outlined tiles, while broken/fragmented LINE-1 copies are shown as grey tiles.

To visualize this structure, we inferred relationships among all LINE-1 units from the chromosome 11 locus, assigned related units to sequence groups, and used those group labels to represent array composition across all 94 haplotypes. In this display, each row corresponds to a haplotype and each column to an ordered LINE-1 position within the array. Coloring units by phylogenetic group shows how copy number and sequence composition vary together across haplotypes. Strikingly, each haplotype contained LINE-1 units from one of the major groups of repeat units (Figure 3, left or right) but not a mixture of both. This representation reveals that the locus is not simply variable in length; it is variable in the specific combinations and arrangements of related LINE-1 units that each haplotype carries.

### Neighbor-distance profiles reveal the tempo of recent human LINE-1 activity

Classical models of LINE-1 evolution describe a ladder-like succession of subfamilies, in which one dominant active group gives rise to the next. The model is powerful and supported by prior phylogenetic and subfamily-based reconstructions of LINE-1 history, but it is necessarily based on the sequence information that remains after many LINE-1 copies have aged. Older elements accumulate substitutions, truncations, internal deletions, and other forms of decay, obscuring the fine-scale relationships that once connected source elements to their descendants. Young full-length LINE-1s provide a different kind of record. Because they have had less time to decay, the youngest LINE-1s remain close in sequence space to their mothers and, sometimes, source-informative flanking sequence. These features allow recent source-element dynamics to be studied at a resolution that is not available from older LINE-1 families. One key advantage of the HPRC assemblies is not simply that they contain more LINE-1s, but that they preserve the young, full-length, allele-resolved sequence relationships needed to reconstruct recent evolution before those relationships are erased.

A distinctive feature of retrotransposon evolution is that each successful element leaves behind molecular copies of itself. These copies are not only products of activity; they are also historical records of the sequence state of successful source elements. Therefore, the local density of related LINE-1 sequences in sequence space can be interpreted as a proxy for past reproductive success. If many elements cluster closely together, at least one element or closely related group of elements in that neighborhood likely generated many daughters over a relatively short evolutionary interval. Conversely, isolated elements or elements with only distant neighbors are less likely to represent recent high-output source lineages. In this sense, full-length LINE-1 sequence neighborhoods provide an evolutionary readout of source-element success, even when the original source element is no longer uniquely identifiable.

To resolve recent LINE-1 activity from pangenome-scale sequence variation, we summarized each of 2,267 full-length LINE-1 consensus sequences by its neighbor-distance profile: the number of other full-length LINE-1s found across bins of pairwise substitutions per kilobase. Elements with many very close neighbors are expected to reflect recent or ongoing retrotransposition, whereas elements whose nearest relatives are more distant represent older expansion phases. This distance-profile approach therefore provides a relative activity clock for human LINE-1s, complementing classical source-element and subfamily analyses [19,20,28].

Gaussian mixture modeling (GMM) of these profiles separated the HPRC LINE-1 set into 12 ordered activity clusters (Figure 4A). The clusters form a strong tempo gradient: cluster 1 has a median neighbor distance of approximately 1.4 substitutions/kb, while cluster 12 has a median of approximately 52 substitutions/kb. The youngest clusters are enriched for the most recently active human LINE-1 lineages: clusters 1-3 contain 426 elements and are dominated by Ta1d LINE-1s (363/426; 85%). In contrast, the oldest clusters are overwhelmingly L1PA2: clusters 7-12 contain 1,379 elements, of which 1,356 (98%) are L1PA2. Between these poles, cluster 4 is dominated by Ta0 but remains mixed with Ta1nd, nonTa, and Ta1d elements, while clusters 5-6 are predominantly nonTa. This ordered structure recapitulates the known succession of human LINE-1 subfamilies while resolving it as a quantitative continuum of recent expansion activity [12,28,41].

**Figure 4.**
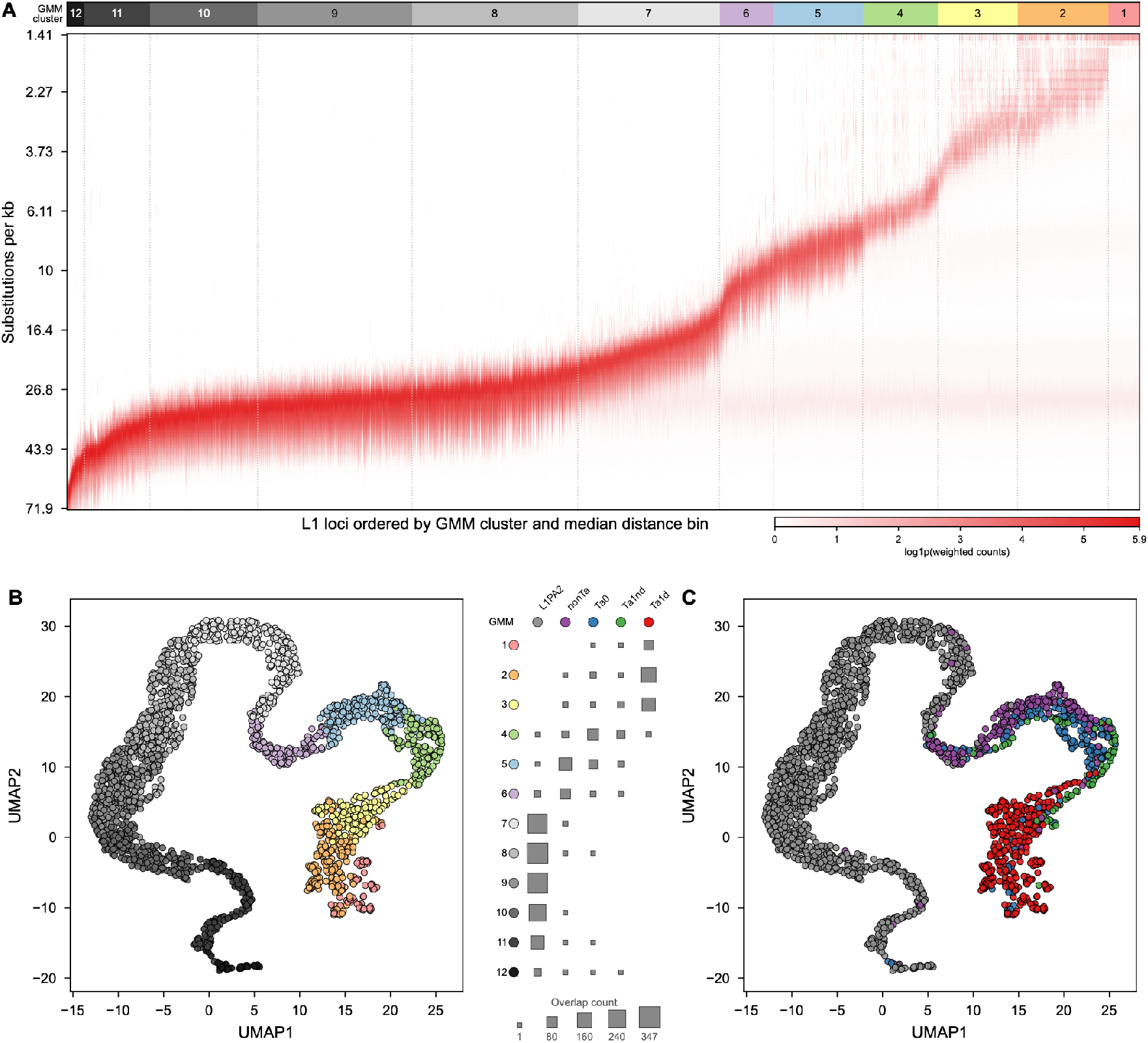
Activity-cluster structure of full-length LINE-1 loci under an ordered cluster palette. (A) Heat map of neighbor-distance profiles for 2,267 full-length LINE-1 loci, with loci ordered by GMM cluster and median distance bin. The grayscale-to-pastel cluster strip emphasizes relative progression across activity-defined groups, while heat-map intensity shows log1p-transformed weighted neighbor counts. (B) UMAP projection colored by ordered GMM cluster. (C) The same UMAP projection colored by LINE-1 subfamily. The central overlap summary shows how activity-defined clusters partition across L1PA2, nonTa, Ta0, Ta1nd, and Ta1d annotations. This view supports the manuscript narrative that recent LINE-1 activity groups broadly track subfamily history while retaining structure not captured by subfamily labels alone.

### Activity clusters broadly track subfamilies but reveal additional structure

The UMAP projection of the same neighbor-distance profiles confirmed that the GMM clusters reflect coherent structure in activity space rather than a one-dimensional sorting artifact (Figure 4B). Young Ta1d-rich clusters occupy the close-neighbor end of the manifold, older L1PA2-rich clusters occupy the oppo site end, and intermediate clusters bridge these extremes. Thus, recent human LINE-1 history is not simply a binary contrast between active and inactive elements, but a succession of partially overlapping expansion waves with different degrees of residual sequence proximity.

Comparing the activity clusters with canonical LINE-1 subfamily annotations showed that subfamily identity explains much, but not all, of the activity structure (Figure 4C). The strongest examples match expectation: cluster 1 is 94% Ta1d, clusters 7-11 are nearly all L1PA2, and cluster 12 remains mostly L1PA2 despite its small size. However, several transition clusters are compositionally mixed, especially cluster 4, which contains Ta0, Ta1nd, nonTa, Ta1d, and one L1PA2 element. This mismatch indicates that neighbor-distance profiles capture recent expansion behavior that is related to, but not redundant with, subfamily labels.

Together, Figure 4 reframes recent human LINE-1 activity as a tempo of lineage turnover. Young Ta1d-rich groups carry the strongest signature of recent expansion, older L1PA2 groups record deeper amplification history, and mixed intermediate clusters mark transitions between these phases. This framework sets up the next question: which sequence changes or source-element properties allowed particular LINE-1 lineages to rise, expand, and then be replaced by their descendants.

### Coarse lineage networks recover directional LINE-1 turnover

Building on the activity tempo defined above, we next asked which nucleotide changes accompany transitions between recent human LINE-1 lineages. We first represented 2,232 full-length non-recombinant LINE-1 consensus sequences as an RMST (randomized minimum spanning tree) [42] built from pairwise sequence distances. We then used recursive tree-bottleneck partitioning to identify high-contrast edges in this graph, yielding 37 lineage neighborhoods connected by 36 split edges (Figure 5A). The graph was oriented from the older end of the recent LINE-1 spectrum, using the neighborhood most enriched for the oldest Figure 4 activity class as the root. This provides an older-to-younger ordering across the recent L1PA2/L1HS continuum, while preserving the important caveat that graph nodes are observed LINE-1 sequences rather than reconstructed ancestors. Edge labels therefore mark representative split-defining substitutions, not an exhaustive catalog of mutations or proof of causality.

**Figure 5.**
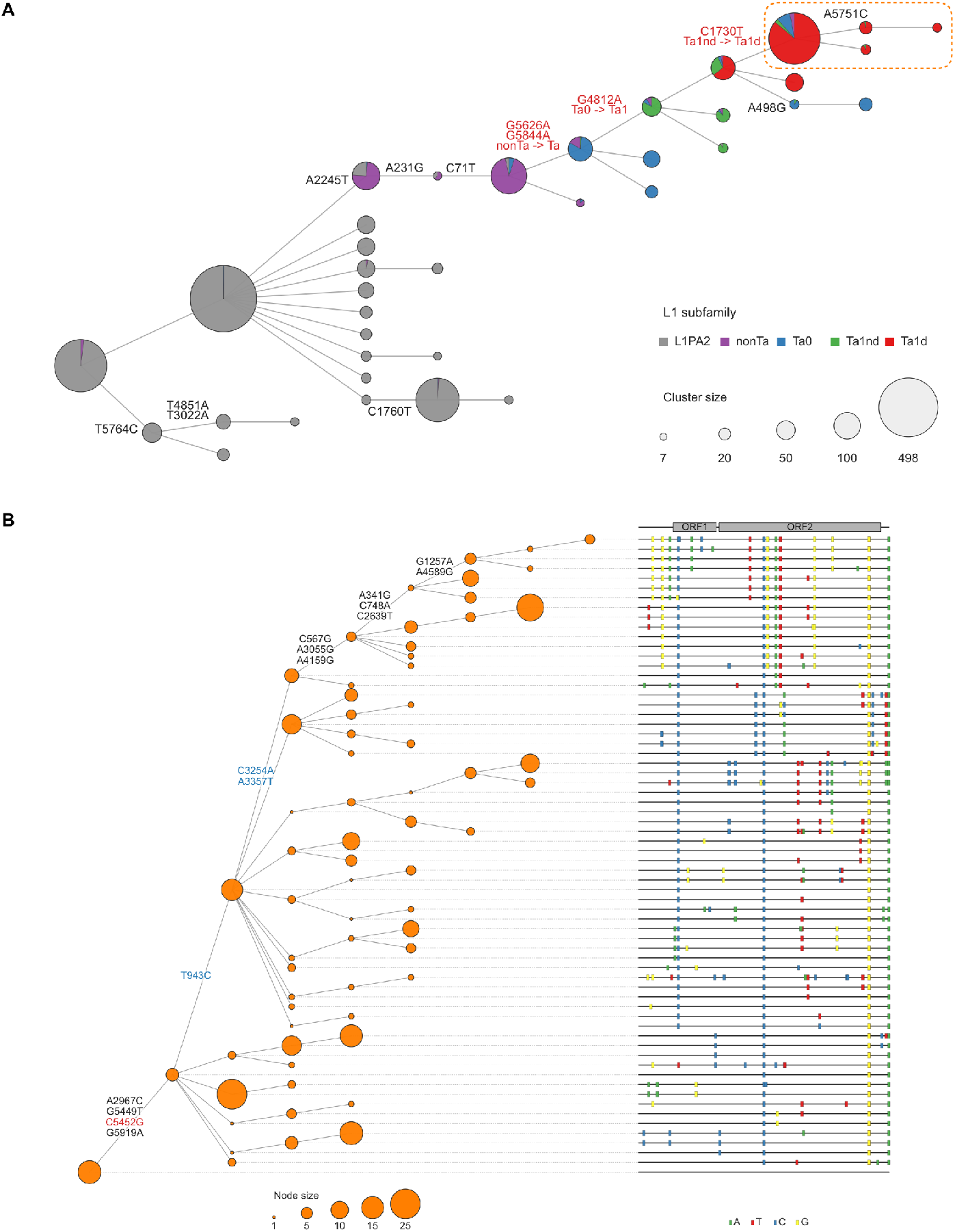
Sequence-defined LINE-1 lineages and nucleotide transitions marking recent LINE-1 diversification. (A) Network-level partitioning of full-length LINE-1 sequence diversity into major lineages. Full-length LINE-1 sequences were organized on an RMST-based graph and recursively split at high-contrast edges to identify coherent local sequence neighborhoods while avoiding partitions driven by single rare tips. Nodes summarize the resulting clusters; node size scales with membership and pie sectors show LINE-1 subfamily composition. Edge labels indicate representative nucleotide changes that best separate the two sides of a split. Red annotations mark transitions that coincide with changes in dominant LINE-1 subfamily, highlighting key lineage transitions along the recent LINE-1 trajectory. (B) Detailed view of the highlighted young lineage from panel A. A rooted row-tree displays the refined young-lineage network together with representative full-length sequence differences relative to the root representative sequence. Orange node size scales with membership, rows are aligned to the full-length LINE-1 sequence track, ORF1 and ORF2 are shown above the track, and colored markers indicate the base observed at each plotted difference site.

The lineage scaffold recapitulates the known succession of human LINE-1 subfamilies while placing specific sequence changes onto the branches that separate them (Figure 5A). Older neighborhoods are dominated by L1PA2 elements, whereas one young branch is strongly enriched for Ta1d elements and for the youngest activity classes defined in Figure 4. Several displayed substitutions coincide with classical subfamily transitions: G5626A and G5844A mark a nonTa-to-Ta branch, G4812A marks a Ta0-to-Ta1 branch, and T1730C marks a Ta1nd-to-Ta1d branch. These transitions extend earlier subfamily-based reconstructions of recent human LINE-1 evolution by placing diagnostic and candidate lineage-defining sites into a pangenome-scale sequence graph [12,28].

### Fine-scale young-lineage analysis resolves mother–daughter relationships among individual loci

We then resolved the young Ta1d-rich branch at higher sequence resolution (Figure 5B). This zoomed analysis includes 335 LINE-1s, of which 292 are annotated as Ta1d, with smaller contributions from Ta0, Ta1nd, and nonTa elements. These LINE-1s are concentrated in the youngest activity classes, with 330 of 335 assigned to the three youngest Figure 4 groups. Using a refined CpG-aware network workflow, 324 core LINE-1s were used to build the backbone and 11 additional LINE-1s were assigned back to nearby nodes, producing a 66-node, 78-edge display. The row-tree layout is paired with a 5,930-bp full-length alignment and 621 representative differences relative to the root/reference row, allowing candidate lineage changes to be read in the context of ORF1, ORF2, and the surrounding UTR sequence.

Several of the strongest Figure 5B transitions fall at sites previously implicated in recent Ta diversification. One early branch in the zoomed tree carries G5449T and C5452G, the two Ta1 hallmark changes separating Ta0-like from Ta1-like states. Additional backbone labels include C943T, A3254C, and T3357A, corresponding to the Chuang/Devine Ta1d diagnostic positions 1026, 3337, and 3440 that define Ta1d-CAT, Ta1d-CCA, and Ta1d-TCA subgroups [43]. In the 335-LINE-1 Figure 5B input, all three sites remain polymorphic, and their placement on different branches indicates that young Ta1d diversification is not captured by a single diagnostic step. Thus, Figure 5B resolves the youngest expansion signal from Figure 4 into nested sequence transitions within the currently most active human LINE-1 lineage.

Together, these results show that young LINE-1s retain a uniquely detailed record of recent retrotransposon evolution. HPRC assemblies make it possible to move from individual intact LINE-1 repertoires, to population distribution, to structurally complex loci, to activity clusters, to lineage transitions, and finally to specific mother–daughter relationships among young loci. This hierarchy transforms the study of human LINE-1 evolution from a coarse subfamily framework into an allele-resolved account of how source lineages rise, diversify, and leave descendants in modern genomes.

## Discussion

Pangenome resolution changes the scale at which recent human LINE-1 evolution can be observed. Single references and short-read population catalogs established that LINE-1 insertions are polymorphic and informative for human variation, but they were limited in their ability to recover full-length, highly similar, recently active copies with haplotype context [15,17,18]. The combination of T2T-CHM13/hs1 and phased HPRC assemblies provides a different view, showing young LINE-1s can be resolved as full-length alleles from individual haplotypes [23,24]. Importantly, these young LINE-1s preserve resolution of evolutionary changes that older copies have largely lost. Deep LINE-1 histories remain visible as coarse subfamily transitions, but mutation, deletion, truncation, gene conversion, recombination, and sequence decay erase much of the fine structure needed to reconstruct recent source-lineage relationships. By contrast, young full-length LINE-1s retain small sequence differences, near-neighbor relationships, intactness states, and population-frequency patterns, allowing recent LINE-1 evolution to be studied as allele-resolved lineage dynamics, expansion tempo, and adaptive changes.

Using these pangenome data, we resolved 2,275 genomic sites harboring full-length LINE-1s across 94 HPRC haplotypes, including 683 sites with an intact allele in at least one haplotype. These intact-LINE-1-containing sites accounted for 13,617 LINE-1 alleles with intact ORF1 and ORF2 coding potential. Across individuals, the diploid intact LINE-1 repertoire centers near 290 elements per genome, closely matching the approximately 300 intact LINE-1s expected from combining CHM1 and CHM13 haploid references [21]. However, similar counts do not mean identical repertoires. Each haplotype carries a high-frequency core of intact alleles shared by many people and a tail of low- and intermediate-frequency intact alleles. These common intact LINE-1s and rare intact LINE-1s answer different biological questions. Common intact alleles define a shared source-element reservoir inherited across many haplotypes, whereas rare alleles mark the edge of current discovery and may be enriched for recent, population-structured, or incompletely sampled variation. Frequency alone does not define activity, but it provides the population context needed to interpret candidate source elements.

The singleton-rich frequency spectrum and continued rise of singleton and low frequency LINE-1s with each additional long-read genome demonstrates that the population-scale reservoir of intact LINE-1 alleles is still highly undersampled. A simple Watterson-style discovery model and singleton/doubleton lower-bound estimates place the broader human pool of segregating intact LINE-1s on the order of several thousand rather than hundreds. However, these estimates depend on sampling design, ancestry representation, assembly sensitivity, and how difficult repeat-rich regions are treated. Similar analyses with high resolution genomes from numerous trios will be important to establish the distribution of rates of de novo germline LINE-1 insertions across populations, and ultimately a more accurate estimate of the total number of potentially active LINE-1s in the human population.

The three-class allele-state matrix adds a second layer to this population view by separating insertion presence from intact coding potential. Earlier MEI catalogs established that LINE-1 insertion profiles track human population structure [15,17,18], and our data confirms that allele-resolved LINE-1 profiles retain this information (Figure 2). However, the HPRC matrix also distinguishes absence from interrupted presence and intact presence. This distinction converts population distribution into an evolutionary signal where insertion presence reflects ancestry, drift, and inheritance; intactness reflects the possibility of retrotransposition competence; and the combination reveals how LINE-1 insertions can persist while their coding potential changes over time. For example, many LINE-1 insertions are present in nearly all haplotypes but intact in only a subset, whereas a smaller set remains both common and intact. Thus, high-frequency LINE-1s are not a single biological class, as they include old insertion sites undergoing molecular decay and broadly shared intact alleles that have preserved ORF1 and ORF2 coding potential across many descendant haplotypes.

The chromosome 11 LINE-1 array extends this logic from insertion-state variation to local structural variation. Most of our analyses treat LINE-1s as retrotransposition-created insertion sites, but the chromosome 11 locus shows that LINE-1 copy number can also change through local duplication, deletion, and rearrangement. This locus contains conserved flanking anchors and a hypervariable internal array whose LINE-1 copy number ranges from 0 to 15 across haplotypes. The array is not simply longer or shorter across genomes; it contains related but sequence-distinct LINE-1 units organized into haplotype-specific combinations. This provides a local, high-resolution example of why haplotype-resolved assemblies are essential for LINE-1 studies. A single-reference or short-read view would likely collapse the locus into a simplified representation, while the HPRC assemblies reveal variable array length, repeat-unit identity, and haplotype-specific organization.

The chromosome 11 locus also suggests that some structurally amplified LINE-1-derived sequences could retain biological relevance even when they are not autonomous source elements. Many copies in the array are not intact by the ORF1/ORF2 definition used for source-element candidates but retain a 5’ UTR and intact ORF1p-coding region. These copies should therefore not be treated simply as inert repeat fragments. They occupy an intermediate category of structurally amplified LINE-1-derived sequences with preserved regulatory and ORF1-associated features that could influence local transcriptional regulation, recombination, or LINE-1 biology.

The neighbor-distance analysis leverages the self-replicating nature of retrotransposons. When an element or closely related group of elements is successful, it leaves daughters that remain close in sequence space. Therefore, the local density of related full-length LINE-1 sequences can be used as a relative readout of past source-element fitness. This does not identify a single mother for every daughter, and it does not prove that all nearby copies came from one source. But it provides a practical activity metric in which close-neighbor-rich groups mark recent expansion phases and distant-neighbor groups mark older amplification history. This approach resolves recent human LINE-1 evolution as a tempo of partially overlapping waves (Figure 4). Ta1d-rich clusters occupy the closest-neighbor end of the distribution, L1PA2-rich clusters occupy the older end, and mixed clusters mark transitional regions between these phases.

This tempo refines, rather than replaces, the classical ladder model of LINE-1 evolution. The canonical model is useful because recent human LINE-1s do show directional turnover where older subfamilies give rise to younger ones, and diagnostic substitutions mark broad transitions. Our data preserve that backbone but also reveal shorter-timescale structure. Activity-defined clusters broadly track subfamily labels, yet they do not map perfectly onto them. Some subfamilies are distributed across multiple activity classes, and several activity classes contain mixtures of Ta0, Ta1nd, nonTa, and Ta1d elements. The simplest interpretation is that the ladder model is a low-resolution projection of a more branched recent process. A dominant directional path is visible, but each rung contains multiple source groups, local expansions, and alternative sequence routes. The young elements reveal this structure because they have not yet accumulated enough mutation and decay to erase their near-neighbor relationships.

At the coarse scale, the RMST/tree-bottleneck scaffold recovers broad lineage neighborhoods and places representative substitutions on the branches separating them (Figure 5). This view recapitulates known transitions such as nonTa-to-Ta, Ta0-to-Ta1, and Ta1nd-to-Ta1d. At the fine scale, the young Ta1d-rich branch can be resolved into individual loci and local sequence neighborhoods, approaching the level of mother-daughter relationships. This is the level of resolution that is largely unavailable for older LINE-1 families. Instead of seeing only subfamily labels, we can ask how specific daughters differ from their close relatives and whether candidate changes mark the origin of a local expansion, accumulate among descendants, or distinguish one daughter branch from another.

The sequence changes that define these transitions (Figure 5) should be interpreted as candidate lineage markers and functional hypotheses, not proven adaptive mutations. Some substitutions may increase intrinsic retrotransposition capacity by altering promoter output, ORF1p-RNA assembly, ORF2p enzymatic activity, RNA processing, or 3’ end formation. Others may change how a LINE-1 interacts with host restriction. Host defenses act at many stages of the LINE-1 life cycle, including transcriptional repression, RNA regulation, ribonucleoprotein stability, reverse transcription, and integration. Factors such as APOBEC3 proteins, MOV10, TREX1, ZAP/ZC3HAV1, TEX19/TEX19.1, and chromatin-based pathways provide multiple opportunities for LINE-1 sequence changes to alter susceptibility to restriction [11,45-49]. Thus, a substitution that rises with a successful lineage could be adaptive because it increases biochemical activity, reduces recognition by a restriction factor, compensates for a host-imposed cost, or functions only in combination with other changes in the same source-element background.

Several limitations shape how these results should be interpreted. First, HPRC phase 1 is not a balanced global sample. It is purposefully enriched for African and admixed American genomes with fewer European, East Asian, and South Asian assemblies. The population enrichments should therefore be read as broad AFR/non-AFR signals rather than final assignments of population specificity (Figure 2). In addition, consensus sequences and insertion-site grouping simplify allelic diversity within loci. They are necessary for population-scale analysis, but follow-up work should test whether specific allele sequences, not just site-level consensuses, differ in activity. Finally, the candidate substitutions provide little insight into the selection pressures (or drift) that drove their evolution. The next step is to connect this allele-resolved evolutionary map to molecular function.

Together, these results show that recent human LINE-1 evolution can now be studied as an allele-resolved population process rather than only as a coarse succession of subfamilies. Haplotype-resolved pangenomes reveal that individual genomes carry a relatively constrained number of intact LINE-1s but draw from a much larger and still incompletely sampled population reservoir of intact source-element candidates. Allele intactness and presence calls connect this repertoire to population structure and molecular decay; the chromosome 11 locus shows how local structural variation can amplify LINE-1 sequence; neighbor-distance profiles resolve a tempo of recent retrotransposition; and sequence networks connect that tempo to candidate lineage-defining substitutions. The emerging picture is of young LINE-1 lineages that diversify below canonical subfamily labels, expand in partially overlapping waves, and potentially adapt through changes that alter intrinsic activity, interaction with host restriction pathways, or other undiscovered selection pressures.

## Methods

### Data acquisition

We analyzed the phase-1 Human Pangenome Reference Consortium assembly panel, consisting of 47 individuals represented by 94 phased haploid assemblies [23]. Assembly FASTA files and sample metadata were obtained from the public HPRC release resources. Sample sex and population metadata were retained for haplotype-level, diploid-level, and population-stratified analyses. Throughout the manuscript, paternal haplotypes from male individuals were treated as Y-bearing haplotypes, and all other haplotypes were treated as X-bearing haplotypes.

For reference-coordinate harmonization, we used the telomere-to-telomere CHM13 v2.0 human reference assembly, corresponding to UCSC hs1 [24]. The hs1 reference FASTA and RepeatMasker annotation track were obtained from the UCSC Genome Browser. Flanking regions of HPRC LINE-1 insertion sites were lifted over to hs1 coordinates so that homologous insertion sites could be compared across haplotypes and summarized in a shared pangenome matrix.

For element-level annotation and comparison to previously characterized intact LINE-1s, we used L1RP as the ORF1 and ORF2 protein reference sequence and incorporated CHM1/CHM13 intact LINE-1 annotations from our prior haploid-reference analysis [21]. File S1 provides the curated HPRC insertion-site matrix together with HPRC metadata, allele-level LINE-1 records, haplotype- and diploid-level LINE-1 count summaries, and intact-LINE-1 discovery summaries. The full-length LINE-1 consensus alignment used for sequence-distance, activity-tempo, recombination, and lineage analyses is provided as File S2.

### LINE-1 sequence identification and curation

Full-length LINE-1s were identified in each HPRC haploid assembly using the current RepeatMasker-based mode of HapLongLINEr (https://github.com/leiyangly/HapLongLINEr), following the general strategy used in our prior haploid-reference analysis [21]. HapLongLINEr parsed RepeatMasker annotations, collapsed fragmented LINE-1 annotations into locus-level calls when appropriate, and retained young LINE-1 annotations meeting the full-length cutoff of at least 5,000 bp. For the young LINE-1 call set used in downstream pangenome analyses, candidate annotations included L1HS, L1PA2, and L1PA3-class records.

For each candidate LINE-1, HapLongLINEr extracted the corresponding assembly sequence and lifted the locus to hs1 using whole-genome assembly-to-reference alignments generated with minimap2 [44]. Lifted coordinates were used to group homologous insertion sites across haplotypes. Candidate loci with ambiguous placement, likely duplication-based origin, or cases where a young LINE-1 inserted into an older LINE-1 annotation were manually curated so that downstream analyses represented homologous insertion sites rather than annotation artifacts.

Coding potential was assessed from the extracted full-length LINE-1 sequence. Open reading frames were identified with EMBOSS getorf using translated ORFs of at least 270 amino acids, and candidate ORFs were searched with BLASTP against an L1RP-derived ORF1p/ORF2p reference database [45,46]. For each LINE-1, the longest ORF1p and ORF2p alignments were retained. A full-length LINE-1 allele was classified as intact only when ORF1p spanned amino acids 1-338 and ORF2p spanned amino acids 1-1275 of the corresponding L1RP reference proteins; full-length alleles lacking one or both complete ORFs were classified as present but interrupted.

After insertion sites were grouped, all allele sequences assigned to the same site were extracted and aligned with MAFFT [47]. A site-level consensus sequence was generated from each alignment with the EMBOSS [45] cons function. These curated site calls form the insertion-site matrix in File S1, and the resulting full-length consensus alignment forms File S2 for downstream distance, recombination, activity-tempo, and lineage analyses.

### Recombinant detection

Recombinant LINE-1 consensus sequences were identified using a custom OpenRDP-based workflow applied to a trimmed, L1RP-anchored master alignment [48]. After removal of the eight manually flagged outlier loci, 2,267 HPRC LINE-1 consensus sequences were screened. For each query sequence, a broad nomination screen was performed with 3SEQ using the query and representative sequences from stable phylogenetic donor groups. Candidate recombinants were required to resolve to a query-specific pair of donor groups and to pass alignment-quality checks around the inferred breakpoint.

Broad-screen candidates were then evaluated by exhaustive focused BOOTSCAN confirmation. For each nominated donor-group pair, all query-centered triplets were tested, consisting of the query plus one donor sequence from each of the two inferred donor groups. Loci were classified as high-confidence recombinants if a dominant, breakpoint-consistent set of BOOTSCAN triplets supported the nominated donor pair in at least 20% of all donor-pair triplets and included at least three distinct donors from each group. Loci with at least 5% triplet support and at least two donors from each group were classified as probable recombinants; remaining candidates were considered ambiguous. High-confidence and probable recombinants were excluded from downstream Figure 4 analyses, whereas ambiguous candidates were retained. In total, 48 loci were nominated by broad 3SEQ, of which 22 were classified as high-confidence recombinants, 13 as probable recombinants, and 13 as ambiguous, resulting in removal of 35 recombinant calls.

### Visualization of LINE-1 Allele States Across HPRC Haplotypes

Figure 2 was based on the Insertion-site matrix and HPRC metadata worksheets in File S1. We included full-length LINE-1 insertion sites that passed alignment QC and carried an intact allele in at least one HPRC haplotype. For each insertion site and haplotype, allele state was encoded as absent, present but interrupted, or intact. HPRC metadata in File S1 were used to assign each haplotype to individual, sex, subpopulation, and superpopulation. Paternal haplotypes from male individuals were treated as Y-bearing, and all other haplotypes were treated as X-bearing.

Autosomal, X-linked, and Y-linked loci were separated by insertion-site chromosome. The autosomal heatmap contains 648 intact autosomal loci across all 94 haplotypes. The sex-chromosome heatmap contains 34 X-linked loci and one Y-linked locus, with X-bearing haplotypes displayed before Y-bearing haplotypes to make chromosome-specific allele availability explicit. The same color encoding was used for all panels: absent, present but interrupted, and present intact.

Haplotype order was determined from the autosomal matrix so that the primary population structure was learned from loci available across all haplotypes. Each autosomal locus was represented by two binary channels, presence and intactness. Loci with little variation across the cohort were down-weighted, and haplotypes were ordered by average-linkage hierarchical clustering of correlation distances, followed by optimal leaf ordering. The resulting row order was used for the autosomal panel and the AFR/non-AFR enrichment panel. For the sex-chromosome panel, the same order was retained within X-bearing haplotypes, while Y-bearing haplotypes were grouped separately and ordered by population label.

Loci within the autosomal and sex-chromosome panels were ordered by cohort-wide allele-state composition. For each locus, we calculated the fraction of haplotypes in the absent, interrupted, and intact states. These three state frequencies were standardized, reduced to a first principal component, and oriented so that loci with higher presence and interruption but lower intactness were placed toward the older/ high-frequency end of the display. Remaining ties were resolved using presence, interrupted, and intact fractions.

To identify population-enriched loci, we tested autosomal insertion presence between AFR and non-AFR haplotypes. Presence was defined as either interrupted or intact. For each autosomal locus, we compared AFR and non-AFR presence counts using a two-sided Fisher’s exact test and corrected P values across loci using the Benjamini-Hochberg procedure. Loci with FDR <= 0.05 were included in the enrichment panel and ordered by the difference in presence frequency between AFR and non-AFR haplotypes, separating non-AFR-enriched and AFR-enriched sites.

### Identification and Comparative Analysis of Homologous chr11 LINE-1-Rich Regions Across HPRC Haplotypes

We extracted the LINE-1 sequences corresponding to chr11:94,140, 954-94,254,695 from the HPRC phase 1 assemblies. Each haplotype assembly was indexed with samtools faidx and minimap2, and the CHM13 interval was aligned to each assembly using minimap2 -x asm5 -c. For each haplotype, the best-supported homologous interval was selected based on alignment length and number of matching bases. The corresponding target interval was extracted and reverse-complemented when aligned on the minus strand, generating one orientation-normalized homologous chr11 interval per haplotype for downstream analysis.

Each haplotype-specific interval was annotated with RepeatMasker to identify LINE-1 copies and determine their genomic coordinates, strand orientation, and repeat subfamily. RepeatMasker coordinates were then used to extract individual LINE-1 copies as copy-level FASTA sequences. This produced a copy-level dataset containing the LINE-1 elements present within the homologous chr11 interval of each haplotype, enabling comparison of sequence variation, repeat composition, and array organization across the 94 HPRC haplotypes.

For L1HS sequence analysis, copy-level L1HS sequences from all haplotypes were grouped by subgroup and aligned to generate a multiple sequence alignment containing haplotype ID, genomic position, strand, repeat annotation, and copy identity. The main sequence relationship analysis focused on high-confidence near full-length L1HS copies with ungapped aligned length greater than 5600 bp. Identical aligned sequences were collapsed into unique sequence variants, while retaining the number of observed copies represented by each variant. Pairwise distances between unique L1HS variants were calculated directly from the alignment as nucleotide differences, counting base-base differences as substitutions, base-gap differences as indels, and ignoring gap-gap positions.

A minimum spanning tree was inferred from the full pairwise distance matrix using Kruskal’s algorithm with deterministic tie breaking. The MST was rooted at node N072 and split panels were generated from this same MST by removing only the edge between N106 and N090. We chose this split because it preserved the original MST topology while separating the two major sequence groups into more interpretable panels. After removing the N106-N090 edge, the N090-side component was displayed as Panel A and rooted/reference-aligned to N090, whereas the N072-side component was displayed as Panel B and rooted/reference-aligned to N072. In the network, backbone nodes N031, N052, N095, N101, N107, N090, N106, N003, N011, and N072 were used as color seeds, and all descendant nodes inherited the color of the nearest upstream backbone node in the rooted MST. To emphasize shared evolutionary changes, aligned node sequences were compared against component-specific reference nodes, variable alignment positions were identified, and singleton sites, defined as positions where only one node differed from the reference, were removed from the final shared-variation analysis.

To evaluate LINE-1 array composition across haplotypes, all LINE-1 copies detected within the homologous chr11 intervals were compiled, including L1HS and older LINE-1 classes such as L1P1 and L1PA2. Copies within each haplotype were ordered by genomic coordinate to reconstruct the local LINE-1 array structure. Full-length L1HS copies were linked to their sequence-variant groups, while shorter L1HS copies excluded from the full-length network analysis and older LINE-1 classes were retained as separate categories for array-level comparison. Population and superpopulation metadata for each HPRC haplotype were used to compare LINE-1 sequence variation and array organization across genetic ancestry groups.

### Inference of Recent LINE-1 Activity Tempo from Neighbor-Distance Profiles

To infer recent *in vivo* LINE-1 activity from full-length sequence variation, we began with the alignment of 2,275 full-length LINE-1 consensus sequences generated from hs1-lifted insertion sites (File S2). Pairwise sequence distances were calculated from this alignment using the Kimura two-parameter model, with distances scaled as substitutions per kilobase. Before constructing activity profiles, eight manually flagged QC outlier consensus sequences were removed from both rows and columns of the distance matrix because they showed atypical alignment or distance-profile behavior. This yielded a curated set of 2,267 full-length LINE-1 consensus sequences for the activity analysis.

Each curated LINE-1 was then represented by a neighbor-distance profile, defined as the distribution of its distances to all other curated full-length LINE-1 consensus sequences. Self-comparisons were excluded, so each profile summarized distances to 2,266 other elements. Exact zero-distance comparisons were counted separately, and non-zero distances were assigned to 200 global distance bins defined on a log-transformed scale from the 0.1st to 99.9th percentile of all non-zero pairwise distances. Distances below or above this range were assigned to the terminal bins. These binned profiles quantify the local density of related LINE-1 copies around each element: LINE-1s with many close neighbors were interpreted as having stronger evidence of recent *in vivo* expansion, whereas LINE-1s whose nearest neighbors were more distant were interpreted as belonging to older expansion phases.

Before clustering, each neighbor-distance profile was processed to emphasize the informative close-neighbor portion of the distribution while reducing sparse long-distance noise. For each row, the profile was smoothed with a five-bin moving average, the first local maximum was identified, and the first local minimum after that peak was used as a row-specific transition point. Bins beyond this point were down-weighted with a logistic function rather than removed entirely, preserving a small contribution from the distant tail. The resulting profiles were converted to row-wise cumulative distribution functions, standardized across LINE-1s, and reduced by principal component analysis using the smallest number of components explaining at least 99% of the variance.

Gaussian mixture models were fit to the transformed profile matrix across candidate component numbers from 2 to 20. Models used diagonal covariance, ten random initializations, and Bayesian information criterion evaluation; the final component number was selected by a BIC-elbow criterion. This procedure selected 12 activity clusters. Cluster labels were ordered by increasing weighted median neighbor-distance bin, so cluster 1 represents the strongest close-neighbor, recently expanded profile class, while cluster 12 represents the most distant-neighbor profile class. For visualization, the same neighbor-distance profiles were also embedded with UMAP after row normalization and fourth-root transformation; this projection was used only to display the activity space, not to assign GMM clusters. Subfamily annotations were then joined to the clustered LINE-1 set to summarize the overlap between activity clusters and L1PA2, nonTa, Ta0, Ta1nd, and Ta1d labels.

### RMST Scaffold of Recent Human LINE-1 Lineages

To resolve major sequence-defined lineages among recent human LINE-1s, we represented 2,232 full-length non-recombinant LINE-1 consensus sequences as a randomized minimum spanning tree (RMST) built from pairwise sequence distances, an approach used for haplotype-network construction when closely related sequences can support alternative minimum-spanning connections [42]. We then partitioned the RMST using a custom recursive tree-bottleneck procedure, following the general MST-clustering principle that clusters can be represented as subtrees separated by high-contrast tree edges [49]. For each eligible edge, we scored the split by the edge length relative to the median internal edge lengths of the two resulting daughter neighborhoods, with an additional penalty for highly unbalanced splits. This yielded 37 lineage neighborhoods connected by 36 split edges (Figure 5A).

### Identification of lineage-defining nucleotide transitions

After the RMST and recursive tree-bottleneck analysis defined major sequence neighborhoods, we used the resulting graph as a framework for identifying nucleotide changes associated with lineage transitions. The starting point was the full-length consensus alignment in File S2, restricted to the 2,232 non-recombinant LINE-1 consensus sequences used for the lineage graph. Each accepted split edge partitions the graph into two neighboring sets of LINE-1s, allowing the two sides of the split to be compared directly across the full alignment.

For each split edge, we scored alignment positions by comparing nucleotide frequencies on the two sides of the split. Candidate markers were prioritized when one nucleotide state was strongly enriched on one side, coherent within that side, and clearly separated from alternative high-scoring positions. Ambiguous, missing, or gap-dominated positions were not used as representative markers. The plotted edge labels therefore represent selected lineage-defining substitutions rather than an exhaustive catalog of all sequence differences between the two groups. Subfamily annotations were used only after marker selection to interpret whether a labeled transition coincided with known changes among recent human LINE-1 subfamilies, including nonTa-to-Ta, Ta0-to-Ta1, and Ta1nd-to-Ta1d transitions.

To examine the youngest Ta1d-rich branch at higher resolution, we performed a second, branch-focused analysis on the 335 LINE-1s belonging to that lineage. We first identified informative positions within the branch and used a CpG-aware, gap-agnostic representation so that recurrent CpG-context changes and missing alignment states did not dominate the local topology. A backbone graph was constructed from 324 core LINE-1s with limited missingness at informative sites. Eleven additional LINE-1s with higher missingness were assigned back to their nearest supported sequence neighborhoods after backbone construction, preserving membership counts without allowing uncertain sequences to reshape the topology.

The final branch-focused display represents the 335 LINE-1s as 66 sequence neighborhoods connected by 78 edges, including a 65-edge backbone. For each neighborhood, representative nucleotide states were compared across a 5,930-bp full-length LINE-1 alignment, and 621 representative difference markers were plotted along the sequence axis. ORF1 and ORF2 coordinates were mapped from L1RP and displayed with the sequence track, allowing candidate lineage-defining substitutions to be interpreted in the context of LINE-1 coding and noncoding regions.

### Use of generative artificial intelligence tools

Generative artificial intelligence tools, including ChatGPT and Codex (OpenAI), were used during data analysis and manuscript development as writing and analytical aids. Codex was used to assist with data-analysis scripting, code review, debugging, and development of scripts used to generate figure panels and associated summary statistics. During manuscript preparation, AI tools assisted with outlining the manuscript, drafting and revising text, harmonizing terminology across sections, generating alternative phrasings, summarizing figure-level interpretations, identifying potential gaps in methods descriptions, and suggesting ways to organize the Results and Discussion. AI assistance was also used interactively to help interpret exploratory analyses and refine conceptual framing, including the relationship between LINE-1 intactness, population frequency, recent activity, and sequence-based evolutionary inference. All outputs from AI tools were critically evaluated by the authors. The authors independently verified all data analyses, numerical values, references, figure interpretations, and inferences, and made all final decisions regarding manuscript content. The authors take full responsibility for the work.

## Acknowledgments

This work was supported by grants from the National Institutes of Health (R35GM142733 and R21AI174130 to RNM). We thank Joseph McDonald and Pranati Dani for technical assistance.

